# High-throughput laboratory evolution and evolutionary constraints in *Escherichia coli*

**DOI:** 10.1101/2020.02.19.956177

**Authors:** Tomoya Maeda, Junichiro Iwasawa, Hazuki Kotani, Natsue Sakata, Masako Kawada, Takaaki Horinouchi, Aki Sakai, Kumi Tanabe, Chikara Furusawa

**Affiliations:** RIKEN Center for Biosystems Dynamics Research; Department of Physics, The University of Tokyo; Universal Biology Institute, The University of Tokyo

**Keywords:** collateral sensitivity, cross resistance, drug resistance, *Escherichia coli*, evolutionary constraints, laboratory evolution, machine learning

## Abstract

Understanding the constraints that shape the evolution of antibiotic resistance is critical for predicting and controlling drug resistance. Despite its importance, however, a systematic investigation for evolutionary constraints is lacking. Here, we performed a high-throughput laboratory evolution of *Escherichia coli* under the addition of 95 antibacterial chemicals and quantified the transcriptome, resistance, and genomic profiles for the evolved strains. Using interpretable machine learning techniques, we analyzed the phenotype-genotype data and identified low dimensional phenotypic states among the evolved strains. Further analysis revealed the underlying biological processes responsible for these distinct states, leading to the identification of novel trade-off relationships associated with drug resistance. We also report a novel constraint that leads to decelerated evolution. These findings bridge the genotypic, gene expression, and drug resistance space and lead to a better understanding of evolutionary constraints for antibiotic resistance.

## INTRODUCTION

The emergence of antibiotic resistance and multidrug-resistant bacteria is a growing global health concern (May, 2014; O’Neill, 2016; Suk et al., 2019; Zhen et al., 2019), and methodology to suppress the emergence of resistant bacteria is desired. Various mechanisms for antibiotic resistance have been identified, including the activation of efflux pumps, modifications of specific drug targets, and shifts in metabolic activities (Harder et al., 1981; Okusu et al., 1996; Palmer and Kishony, 2014; Toprak et al., 2011; Yelin and Kishony, 2018; Zampieri et al., 2017). Quantitative studies of resistance evolution showed that these mechanisms for resistance are tightly interconnected, as demonstrated by the complicated networks of cross-resistance and collateral sensitivity among drugs (Barbosa et al., 2017; Chevereau et al., 2015; Girgis et al., 2009; Lázár et al., 2014; Suzuki et al., 2014), which is the phenomena where the acquisition of resistance to a certain drug is accompanied by resistance or sensitivity to another drug. Such interactions among resistance mechanisms result in constraints on accessible phenotypes in evolution, which could provide a basis to predict and control the resistance evolution (Furusawa et al., 2018; Imamovic et al., 2018; Lässig et al., 2017). For example, the cyclic or simultaneous use of two drugs with collateral sensitivity, to which pathogens did not easily acquire resistance simultaneously, were demonstrated to suppress resistance evolution (Munck et al., 2014; Yoshida et al., 2017). Thus, elucidating evolutionary constraints are crucial for predicting and controlling the evolution of antibiotic resistance. However, despite its importance, a systematic investigation of evolutionary constraints for antibiotic resistance evolution is still lacking.

Laboratory evolution associated with genotype sequencing and phenotyping is an effective approach to investigate constraints in adaptive evolution (Conrad et al., 2011; Palmer and Kishony, 2013). Here, we performed high-throughput laboratory evolution of *Escherichia coli* under 95 antibacterial chemicals including inhibitors of cell wall synthesis, protein synthesis, DNA replication, RNA polymerase, several metabolic pathways, chelators, heavy metals, etc. (Table S1). Changes in the transcriptome, genomic sequence, and resistance profile in the evolved strains were quantified, resulting in a multiscale dataset for analyzing stress resistance. By applying interpretable machine learning techniques, the emergence of low dimensional phenotypic states in the evolved strains was observed, indicating the existence of evolutionary constraints. The underlying biological processes corresponding to each state were analyzed by introducing the representative mutations to the parent strain. We also report decelerated evolution, where the resistance of the evolved strains in a certain stressor is overtaken by strains evolved in a different stressor. We will show how our resource could provide a quantitative understanding of evolutionary constraints in adaptive evolution, leading to the basis for predicting and controlling antibiotic resistance.

## RESULTS

### Laboratory evolution of *E. coli* under 95 stress conditions

To systematically investigate drug-resistant phenotypes, we performed high-throughput laboratory evolution using an automated culture system (Fig. 1A) (Horinouchi et al., 2014), for 95 stressors covering a wide range of action mechanisms (Fig. 1B, Table S1). To evaluate the reproducibility of the evolutionary dynamics, six independent culture lines were propagated in parallel for each stressor. In total, 576 independent culture series were maintained (95 stressors plus a control without any stressor × six replicates) for 27 daily passages corresponding to about 300 generations. Figure 1A shows examples of the time course of half-maximal inhibitory concentrations (IC_50_s) during laboratory evolution, while all time courses of IC_50_s are shown in Fig. S1. Among the 95 stressors, a significant increase of IC_50_ was observed for 89 stressors (Mann-Whitney U-test, false discovery rate (FDR) < 5%). For further phenotypic and genotypic analyses, we selected 192 evolved strains, i.e., four evolved strains isolated from 47 stressors plus a control without any stress.

**Fig. 1.**
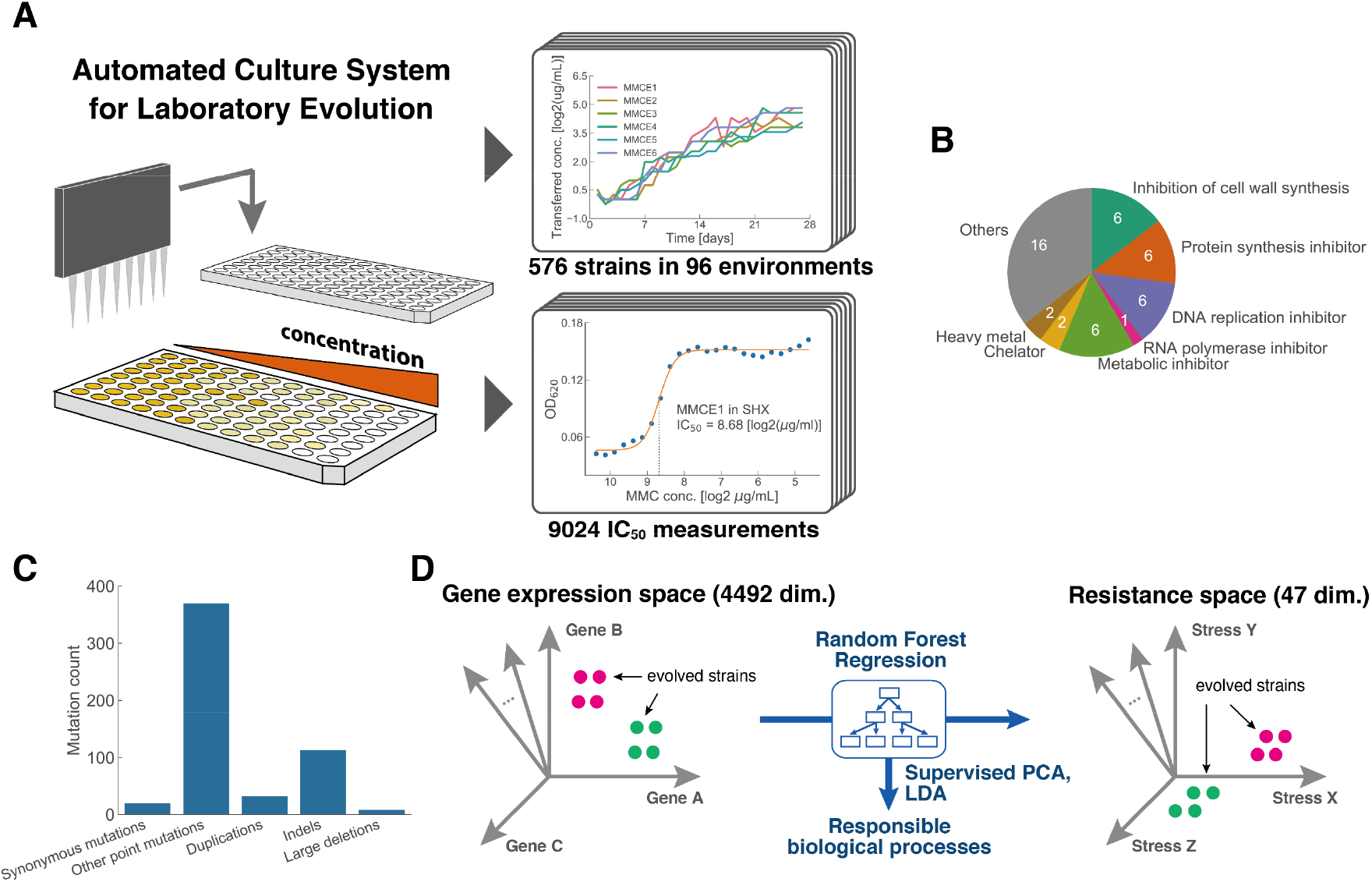
Laboratory evolution of *E. coli* under 95 stress conditions. (A) Schematic of the experimental setup (automated culture system for laboratory evolution). (B) Stress categories for the environments used in the half-maximal inhibitory concentration (IC_50_) measurements. (C) Distribution of mutation events for the evolved strains according to its mutation type, except for strains evolved in mutagens. (D) A random forest regression model was constructed to predict IC_50_ values using the gene expression levels (Methods). Supervised principal component analysis (PCA) was applied to the 213 gene expression levels selected by the random forest algorithm.

### Phenotypic and genotypic changes in evolved strains

To explore phenotypic changes in the 192 evolved strains, we first quantified changes in the stress resistance profiles by measuring the IC_50_ of all 47 chemicals for each evolved strain (9024 measurements in total), relative to the parent strain (Table S2, STAR Methods). The resistance profile measurements allowed us to study how common cross-resistance, a phenomenon where an evolved strain in a certain stress gains resistance to another stress, occurs. By comparing the four evolved replicates and the parent strain, we found that 336 and 157 pairs of stresses exhibited cross-resistance and collateral sensitivity, respectively, within the possible 2162 combinations (Mann-Whitney U-test, FDR < 5%, Fig. S2). Phenotypic changes of the evolved strains were quantified by transcriptome analysis to examine gene expression levels responsible for stress resistance (Table S3, STAR Methods). Details of the expression changes are given in the following sections.

To investigate genetic alterations underlying the observed resistance, we performed genome resequencing analysis of the 192 evolved strains (Table S4). Although some of the evolved strains carried more than 20 mutations, about 80 % of the evolved strains carried less than five mutations. Among the 47 stressors, the highest number of mutations was observed in glutamic acid γ-hydrazide (GAH) evolved strains carrying between 73 to 223 mutations. These numbers were much larger than that of the evolved strains against known mutagens (e.g. 4-nitroquinoline-1-oxide (NQO) and mitomycin C (MMC)), indicating a high mutagenic activity of GAH. Excluding strains evolved with these three mutagens, 21 and 307 mutations were identified as synonymous and nonsynonymous mutations, respectively (Fig. 1C). For strains evolved in non-mutagens, the ratio of nonsynonymous to synonymous mutations per site was 5.26, implying that around 80% of the nonsynonymous mutations were beneficial (Shibai et al., 2017; Tenaillon et al., 2012).

### Supervised PCA reveals modular phenotypic states

To explore the relationship between the gene expression changes and resistance evolution, we performed dimension reduction on the gene expression data using supervised principal component analysis (PCA), which enables the extraction of a subspace in which the dependency between gene expression and stress resistance is maximized (Bair et al, 2006). First, to exclude genes with unchanged or noisy expression in the evolved strains, we used random forest regression to screen the genes which contribute to the prediction of resistance changes (Fig. 1D, see STAR Methods for details), resulting in the selection of 213 genes which had high correlations with resistance levels. Supervised PCA using expressions of these genes revealed the existence of clusters of evolved strains in the dimension reduced gene expression space. To clarify the clusters, we applied hierarchical clustering to the evolved strains (Fig. 2A, see STAR Methods), which demonstrated modular classes of expression profiles. Intriguingly, strains within the same class were not necessarily evolved in the same stress nor stress category. Similar phenotypic convergence of drug resistant strains has recently been reported for *Pseudomonas aeruginosa* (Imamovic et al., 2018). To elucidate characteristic gene expressions for each class, we applied linear discriminant analysis (LDA) (see STAR Methods). This allowed us to extract the most discriminative set of genes for each class, through the observation of each decision boundary (Fig.2B).

**Fig. 2.**
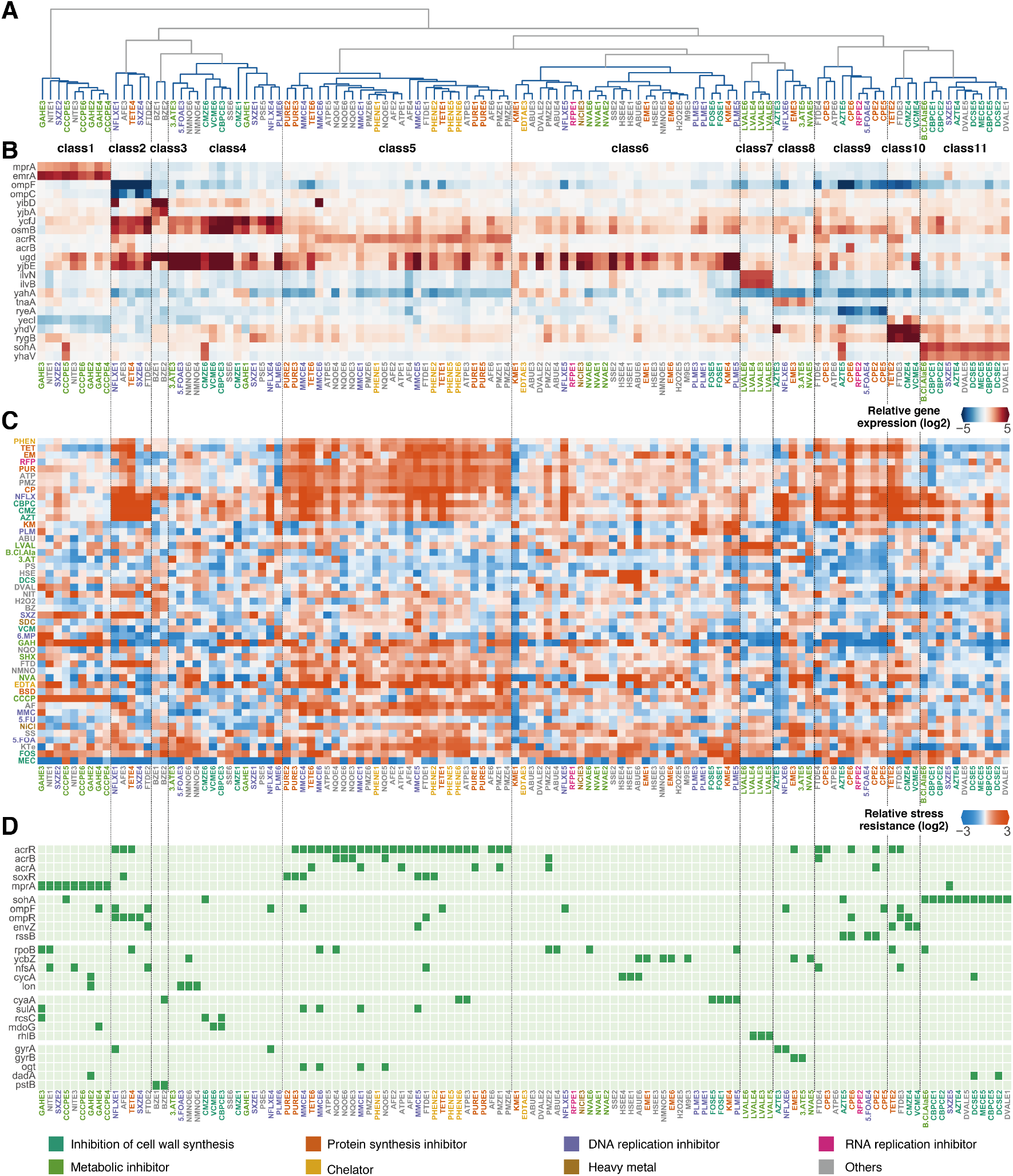
Supervised PCA reveals distinct clusters in the genotype, expression, and the resistance (IC_50_) space. (A) Dendrogram of the result of hierarchical clustering performed in the 36-dimensional supervised PCA space. One cluster and three singletons were omitted due to visibility. The full version is presented in Fig. S3. (B) Gene expression levels of representative genes for each cluster, relative to the parent strain. The genes were selected from the intersection of the top two gene weights for the linear discriminant analysis (LDA) axis and differentially expressed genes (STAR Methods). (C) IC_50_ values relative to the parent strain. Colors for the tick labels correspond to the stress categories. (D) Commonly mutated genes within the evolved strains. Mutated genes enriched for each cluster clarified by Fisher’s exact test (p < 0.01) are presented. Mutated genes that were identified in more than seven strains are also presented. Genes are sorted based on gene ontology categories.

To investigate how the classes of gene expression profiles correspond to stress resistance, we observed the relative IC_50_ for each of the 47 stresses of each evolved strain sorted based on the hierarchical clustering of gene expressions (Fig. 2C). As shown in the figure, the classes in the supervised PCA space correspond well with the stress resistance patterns. To evaluate how accurate the neighboring relationship in the resistance space corresponds to that of the supervised PCA space, we computed the class dissimilarity (*W*_*n*_) in the resistance profile for the classes in the dimension reduced gene expression space (STAR Methods). For comparison, we computed *W*_*n*_ for classes based on the resistance space, the whole gene expression space, and the genotype space (Fig. S3). This metric revealed that clustering in the supervised PCA space coincides well with the resistance space, and better than that of the whole expression space or the genotype space. This indicates that topological relationships between the strains in the resistance space could be accurately represented in a subspace of the gene expression space.

### The relation between mutations and supervised PCA classes

We next wondered how the identified classes in the supervised PCA classes could be characterized. To our surprise, relatively clear relationships between the genome, transcriptome, and resistance profiles were observed for the modular classes in the supervised PCA space. Fig. 2D shows a subset of the commonly mutated genes within the 192 evolved strains. As shown, the patterns of fixed mutations coincide well with the modular classes in the gene expression space, although no genotypic information was used for the hierarchical clustering. This suggests that the identified mutations play a meaningful role in the modular gene expression classes. For example, all evolved strains in class 1 had mutations in *mprA*, which encodes a repressor for multidrug resistance pump EmrAB, while all strains in class 11 had mutations in *sohA*, which encodes the antitoxin for the SohA (PrlF)-YhaV toxin-antitoxin (TA) system.

Although strains in the same class had similar gene expression levels, they did not necessarily share the same mutations. For example, evolved strains in class 5 showed an increased expression of *acrB,* which encodes a component of the AcrAB/TolC multidrug efflux pump, while most of the strains (26/28) in class 5 had mutations in *acrR,* which encodes a repressor for *acrAB* (Fig. 2B, D). Interestingly, the other two strains also showed an increase in *acrB* expression without an *acrR* mutation. Strains in class 8 consistently had an increased expression of *tnaA* encoding tryptophanase, whereas four out of five strains had mutations in genes encoding DNA gyrase subunit A or B (*gyrA* or *gyrB,* Fig. 2B,D). We further confirmed that the introduction of the observed H45Y mutation in *gyrA* to the parent strain leads to an 11.1 ± 6.6-fold increase in *tnaA* mRNA level, through quantitative reverse transcription polymerase chain reaction (qRT-PCR) analysis. Consistent with these results, a previous study also suggested the involvement of a mutation in *gyrB* for increased TnaA production and quinolone resistance (Lee et al., 2010). It should be noted that, NVAE5 which did not have a mutation in *gyrA* nor *gyrB*, also showed an increased expression of *tnaA*. These results suggest the existence of multiple paths in the genotypic space for *E. coli* to reach desired expression and resistance levels.

Commonly decreased expressions of *ompF*, which encodes the outer membrane porin, were observed in several classes (class 2, 9, and 10, Fig. 2D) and were associated with resistance to cell wall inhibitors and other stresses. It is known that a decrease of *ompF* expression could be caused by either inactivation of the OmpR/EnvZ two-component system or RssB, which is a regulator of the alternative sigma factor RpoS (Delcour, 2009; Gibson and Silhavy, 1999; Norioka et al., 1986). Indeed, all strains in class 2 and class 10 had mutations in either *ompR* or *envZ*, and four out of nine strains in class 9 had mutations in *rssB.* Although these strains commonly had a decrease in the *ompF* expression level, they have been classified to different classes due to the consistently different gene expression profiles such as the *ompC* encoding a porin and *rygB* encoding a small RNA involved in the regulation of the outer membrane composition (Fig. 2B). Interestingly, although all three classes showed resistance to β-lactams (e.g., carbenicillin (CBPC), cefmetazole (CMZ)), resistance levels to stresses such as sulfisoxazole (SXZ) and DL-3-hydroxynorvaline (NVA) differed between classes (e.g. strains in class 2 and 10 exhibited resistance to SXZ, while strains in class 9 did not, Fig. 2C, Fig. S2). This suggests that different classes in the supervised PCA space correspond to different stress resistance mechanisms.

Evolved strains derived from the same selection pressure were not always categorized in the same class. For example, none of the four SXZ evolved strains shared the same supervised PCA class (SXZE2 in class 1, SXZE4 in class 2, SXZE1 in class 4, and SXZE5 in class 11), and each strain showed different expression and resistance patterns, indicating a rugged fitness landscape with multiple local peaks. This ruggedness was observed for the four norfloxacin (NFLX) evolved strains as well. Intriguingly, these local peaks were accessible not only by SXZ and NFLX evolved strains, but also by strains which evolved in other stresses (for example, see class 2). These results suggest that evolution under the same selection pressure does not necessarily lead to the same phenotype (Jiao et al., 2016; Nichol et al., 2019), and that these local peaks in the fitness landscape are shared by different stresses. Overall, the low dimensional phenotypic states revealed by our high-throughput measurements loosely corresponded with the genotypic space, suggesting the existence of various genotypic pathways to reach local optima in the fitness landscape, which were shared by strains evolved in diverse stresses.

### Commonly mutated genes provide the basis for chemical resistance

Genome resequencing analysis revealed that several genes were commonly mutated in multiple evolved strains, suggesting the contribution of these mutations to the observed resistance acquisition. The detailed description of these genes and mutations are presented in Table S4. To verify the effects of the commonly mutated genes found in the evolved strains, we introduced 64 of the representative mutations (Table S2) to the parent strain by multiplex automated genome engineering (MAGE) (Nyerges et al., 2016), and quantified changes in the IC_50_s of all 47 chemicals against each strain. We first asked whether the cross-resistance and collateral sensitivities observed within the evolved strains could be reproduced by the 64 reconstructed mutant strains. Accordingly, we calculated the Pearson’s correlation coefficient R between the IC_50_s of all 47 stresses within the evolved strains. For example, evolved strains resistant to CBPC tended to exhibit resistance to aztreonam (AZT) as well (R = 0.95, Fig. 3A). We then calculated correlation coefficients for the reconstructed mutant strains (Fig. 3B) and compared the coefficients with that of the evolved strains. The coefficients for the evolved strains highly correlated with that of the reconstructed mutant strains (R = 0.66, Fig. 3C,D), indicating that the observed collateral relationships within the evolved strains could be explained by the commonly mutated genes. These results allowed us to further investigate the role of mutations in the context of cross-resistance and collateral sensitivities.

**Fig. 3.**
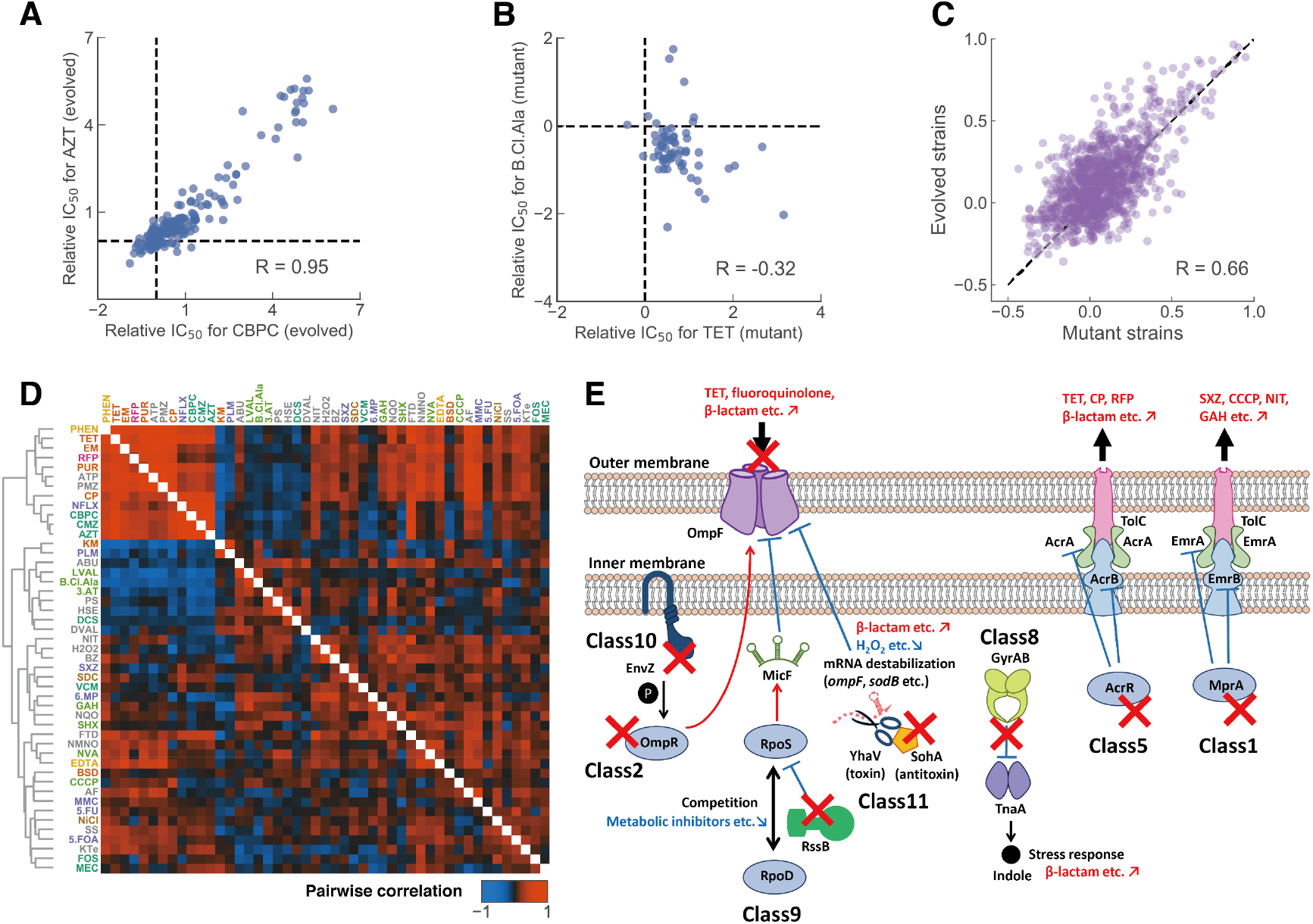
Commonly mutated genes provide the basis for chemical resistance. (A, B) Relationship between the IC_50_ values of carbenicillin/aztreonam (CBPC/AZT) and tetracycline/β-chloro-L-alanine (TET/B.Cl.Ala) for the 192 evolved strains and 64 site directed mutants, respectively. R denotes the Pearson’s correlation coefficient. (C) Relationship between the corresponding pairwise correlation coefficients shown in (D). (D) Pearson’s correlation coefficient for all pairwise combinations of stress resistance for the evolved strains (upper right) and the site-directed mutants (lower left). The order of stresses was determined by hierarchical clustering performed on the pairwise correlation values of the site-directed mutants. (E) Schematic illustration of stress resistance acquisition mechanisms corresponding to the supervised PCA clusters. Typical stresses which exhibited resistance (red) and sensitivity (blue) are shown.

Most of the evolved strains acquired cross-resistance from mutations in genes encoding transporters and porins (Table 1, Table S5). Although it is well known that antibiotic resistance of *E. coli* could be triggered by the overexpression of efflux systems and decreased production of porin proteins (Okusu *et al.,* 1996; Delcour, 2009; Li and Nikaido, 2009), our high-throughput IC_50_ measurements, for both the evolved strains and the mutant strains, allowed us to identify novel substrates for these transporters and porins (Fig. 2D, Table 1, Table S5). For example, activation of the multidrug efflux pump AcrAB/TolC, through the inactivation of its repressor AcrR, resulted in resistance not only to previously described substrates such as tetracycline (TET) and erythromycin (EM) (Ma et al., 1993; Mazzariol et al., 2000; Okusu et al., 1996), which are both known as protein synthesis inhibitors, but also to novel substrates such as NVA (threonine analogue) and NQO (mutagen) (for a detailed list see Table 1). The activation of EmrAB/TolC is regulated by *mprA*, which is also an efflux pump, and resulted in resistance to a previously identified substrate CCCP (uncoupling agent, Lomovskaya et al., 1995), and also novel substrates such as chloramphenicol (CP, protein synthesis inhibitor) and phleomycin (PLM, DNA intercalator) (Fig. 3E, Table S5). We also confirmed that the inactivation of the OmpF porin resulted in resistance to previously described substrates such as CBPC (cell wall synthesis inhibitor) (Balagué and Véscovi, 2001; Harder et al., 1981), and novel substrates such as 1,10-phenanthroline (PHEN, chelator), puromycin (PUR, protein synthesis inhibitor), and other chemicals (Fig. 3E, Table 1). Our results suggest that mutations in the transporters and porins above could lead to resistance to a wide spectrum of drugs with different mechanisms of action. We also identified the contribution of an uncharacterized transporter to chemical resistance. Two D-valine (DVAL) evolved strains had mutations in *yhjE* which encodes a putative transporter. The contribution to resistance was confirmed through the reconstructed *yhjE* inactivation mutant strain which showed more than a 2-fold increased resistance to PLM, NVA, and DVAL (Table S2), which suggests the uptake through YhjE. Overall, our results indicate that the effects of chemical uptake and efflux are major mechanisms for cross-resistance.

**Table 1.**
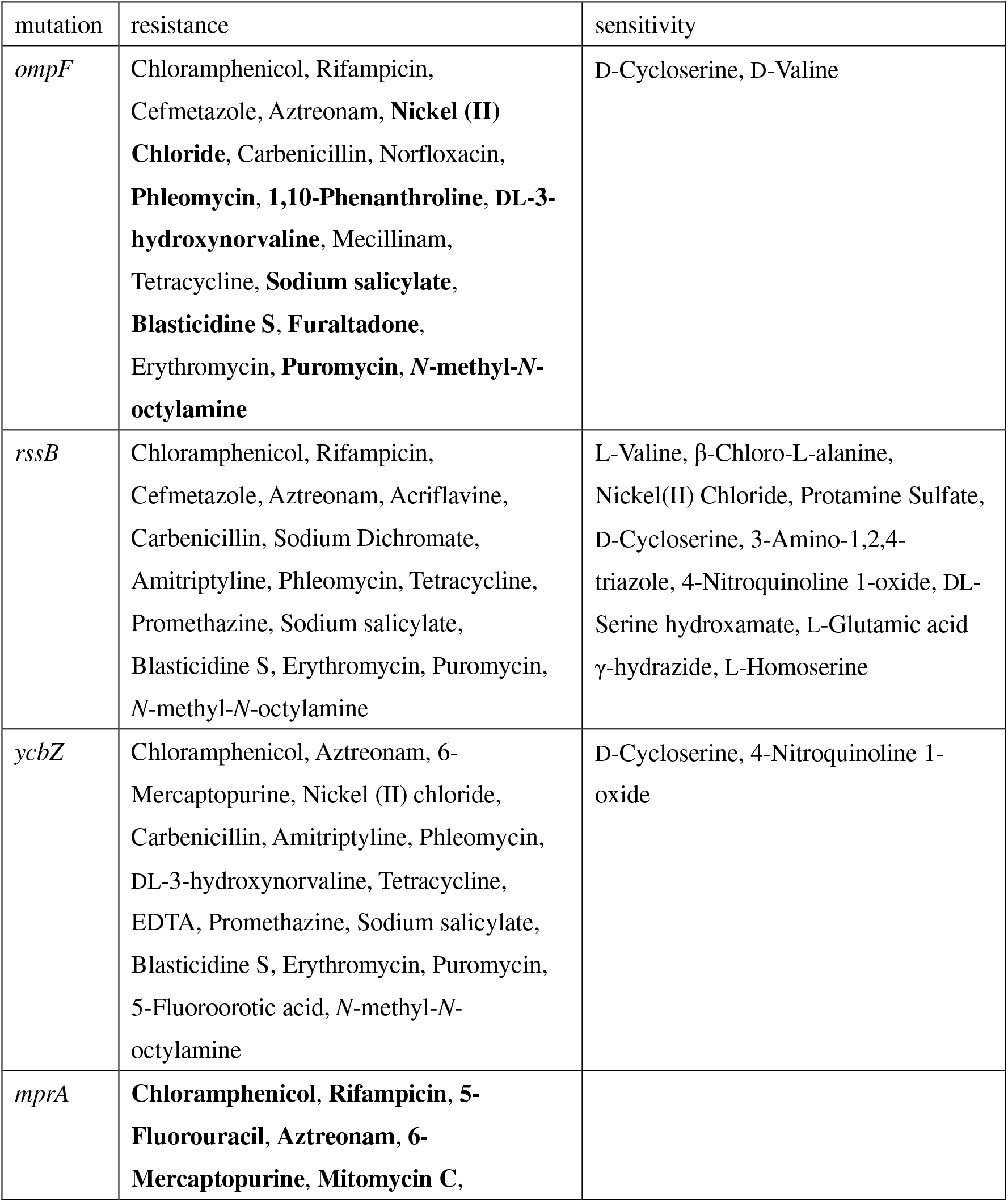
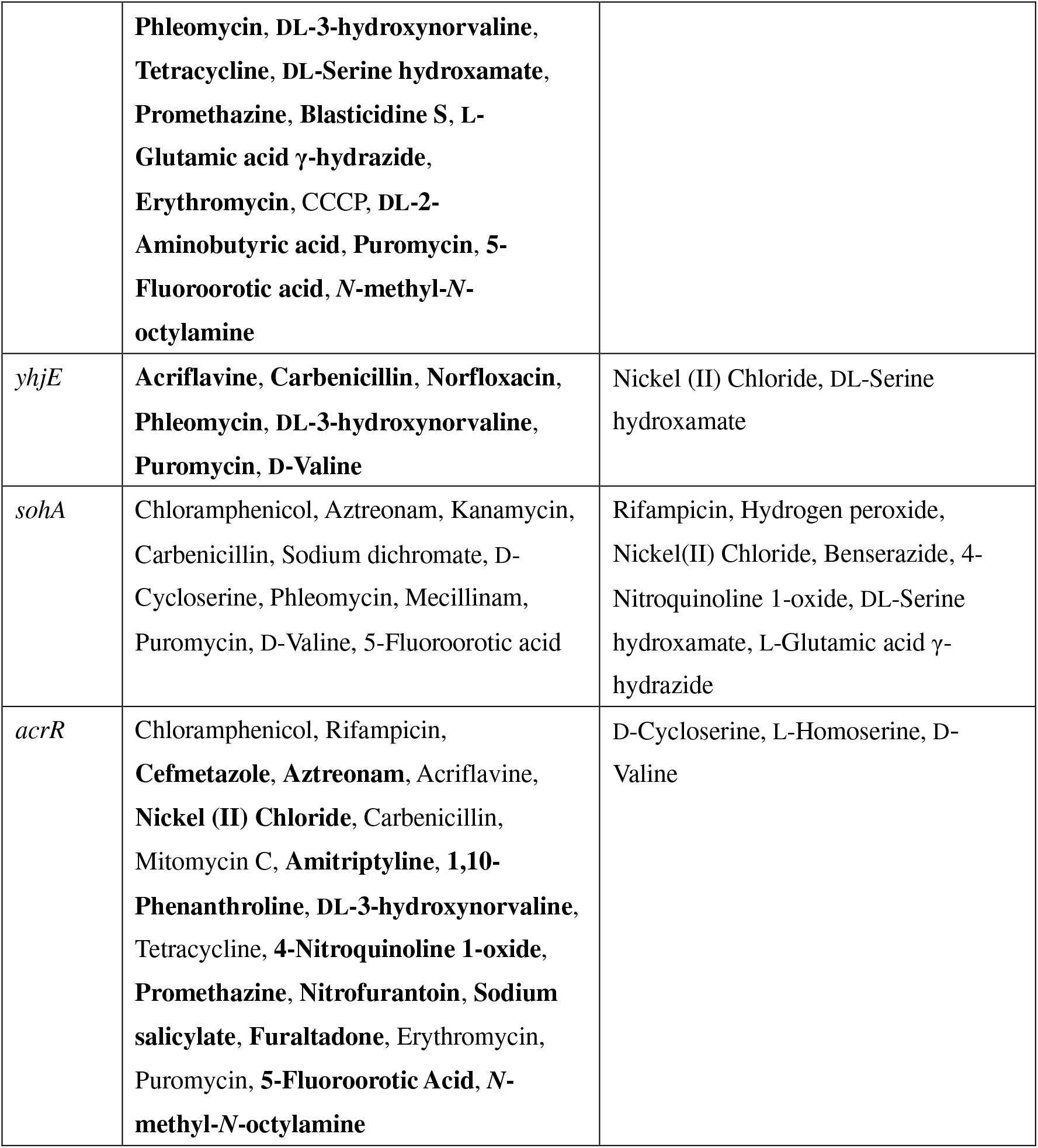
Representative cross-resistances and collateral sensitivities observed in the reconstructed strains. Chemicals that were identified as significantly increased or decreased IC_50_ values (Mann-Whitney U-test, false discovery rate (FDR) < 5%) in the reconstructed *ompF*, *rssB*, *ycbZ*, *mprA*, *yhjE*, *sohA,* and *acrR* mutant strains are shown. For transport machinery (i.e. *ompF*, *mprA*, *yhjE*, and *acrR*), newly identified putative substrates are shown in bold letters. The full list of cross-resistances and collateral sensitivities observed in all 64 reconstructed mutant strains are shown in Table S5.

We have also identified new mechanisms underlying cross-resistance to chemicals. The increased expression of SohA-YhaV TA system along with mutations in *sohA* was observed in class 11 (Fig. 2B, D). Interestingly, all evolved strains carrying the *sohA* mutation, including the 11 strains in class 11, had the same mutation: i.e. duplication of TTCAACA sequences at 272 bp downstream of the start codon (Table S4). Although the contribution of SohA-YhaV to stress resistance has not yet been reported, these evolved strains and the reconstructed *sohA* mutant strain commonly showed resistance to CBPC, AZT, and DVAL (Fig. 2C, Table 1). All 11 strains in class 11 showed a decreased expression of *ompF* (Fig. 2B), and the decrease in *ompF* mRNA level by 0.52 ± 0.01 in the reconstructed *sohA* mutant strain was confirmed by qRT-PCR analysis. These results indicate that cross-resistance to CBPC, AZT, and DVAL by the *sohA* mutation is at least partially caused by decreased expression of *ompF*. Since YhaV is a translation-dependent RNase (Schmidt et al., 2007), this decrease in *ompF* expression might be caused through the alteration of global gene expression.

The contribution of the uncharacterized protein YcbZ to cross-resistance was also confirmed. YcbZ is a putative protease which is shown to be involved in translation and ribosome biogenesis (Gagarinova et al., 2016). We found YcbZ mutations in three EM evolved strains and two N-methyl-N-octylamine (NMNO) evolved strains. These evolved strains and the reconstructed *ycbZ* inactivation mutant strain commonly showed resistance not only to EM and NMNO but also to ethylenediamine-*N,N,N’,N’*-tetraacetic acid, disodium salt, dihydrate (EDTA), NVA, and 5-fluoroorotic acid monohydrate (5.FOA) (Fig. 2C, Table 1).

### Quantification of novel collateral sensitivities

The identified classes in the supervised PCA space revealed novel collateral sensitivity relationships for antibiotic resistance acquisition. For example, the evolved strains in class 9 exhibited sensitivity to metabolic inhibitors such as L-valine (LVAL), β-chloro-L-alanine (B.Cl.Ala), 6-mercaptopurine monohydrate (6.MP), GAH, and 3-amino-1,2,4-triazole (3.AT) (Fig. 2C, Fig. S2). Since 4/9 of the class 9 strains had mutations in *rssB*, we speculated that this observed sensitivity is caused by *rssB*. This was confirmed through IC_50_ measurements of the reconstructed *rssB* inactivation mutant strain which showed a 2 to 5-fold change in sensitivity to the stresses above (Table S2, Fig. S4). Since class 9 strains and the reconstructed *rssB* strain both show resistance to cell wall inhibitors and other stresses (AZT, CBPC, TET), our results indicate a trade-off between these stresses and metabolic inhibitors. It has been reported that while *E. coli* strains with higher *rpoS* levels show increased resistance to several external stresses (Radzikowski et al., 2016), they also show decreased carbon source availabilities and poor competitiveness for low concentrations of nutrients due to the competition between RpoS and the house-keeping sigma factor RpoD (sigma 70) (Ferenci, 2005). Because RssB facilitates the degradation of RpoS, the collateral sensitivities to several metabolic inhibitors in class 9 evolved strains could also be caused by the sigma factor competition.

Collateral sensitivities associated with the newly found *sohA* mediated resistance were also identified. All evolved strains in class 11, all carrying the same *sohA* mutation, exhibited collateral sensitivity to hydrogen peroxide (H_2_O_2_), benserazide (BZ), and NQO (Fig. 2C, Fig. S2). Consequently, the same collateral sensitivities were also observed in the reconstructed *sohA* mutant strain (Fig. S4). These results suggest that DVAL, CBPC resistance, acquired through the *sohA* mutation, leads to a trade-off to H_2_O_2_, BZ, and NQO. Previous studies reported that *E. coli* mutant strains lacking superoxide dismutase showed increased susceptibility to H_2_O_2_ mediated killing (Imlay and Linn, 1987). Indeed, all strains in class 11 consistently showed a 0.45 ± 0.11-fold decrease in *sodB* expression, which encodes (Fe) superoxide dismutase, in the transcriptome data. Through qRT-PCR analysis, the decrease in *sodB* mRNA level by 0.46 ± 0.06 in the reconstructed *sohA* mutant strain was also confirmed. This suggests that the observed H_2_O_2_ sensitivity is caused by the degradation of *sodB* through the YhaV toxin.

### Decelerated evolution against cell wall synthesis inhibitors

In the adaptive evolution to β-lactams (i.e., CMZ and CBPC), we found that some strains which evolved under certain stresses acquired higher resistances to β-lactams than strains which were directly selected by β-lactams. Fig. 4 presents IC_50_ values of CBPC and CMZ observed in the 192 evolved strains, showing that evolved strains under CBPC and CMZ (marked by blue and cyan) did not exhibit the highest resistance levels to β-lactams, but instead, evolved strains under other stresses such as TET and NFLX showed much higher resistance levels (Fig. 4A, B). This “decelerated” resistance evolution to β-lactams was reflected in the difference in the mutation profile among the evolved strains. We found that the evolved strains which exhibited the highest resistances generally had mutations in genes related to the membrane porin protein OmpF, i.e., *ompF*, *ompR* and *envZ* (Fig. 4C). It is known that the disruption of OmpF contributes to resistance acquisition to β-lactams (Harder et al., 1981), and in fact, we confirmed that introduction of the identified *ompF* mutations to the parent strains significantly increased IC_50_ values of CMZ and CBPC (green marker in Fig. 4C). In contrast, the strains evolved under CBPC or CMZ had significantly smaller numbers of mutations in OmpF related genes (one out of eight evolved strains) in comparison with other strains with high β-lactam resistance (p = 0.04). This result suggested that in our setup of laboratory evolution, the fixation of mutations related to OmpF is suppressed under the addition of β-lactams, even though they can increase the resistance to the drug. Possible explanations for this decelerated evolution against β-lactam could be the existence of a fitness cost associated with *ompF* mutation and/or negative epistasis between *ompF* and *sohA* mutations. However, neither such fitness cost nor negative epistasis was observed (STAR Methods, Fig. S5). At present, the mechanism for the decelerated evolution against β-lactams remains unclear.

**Fig. 4.**
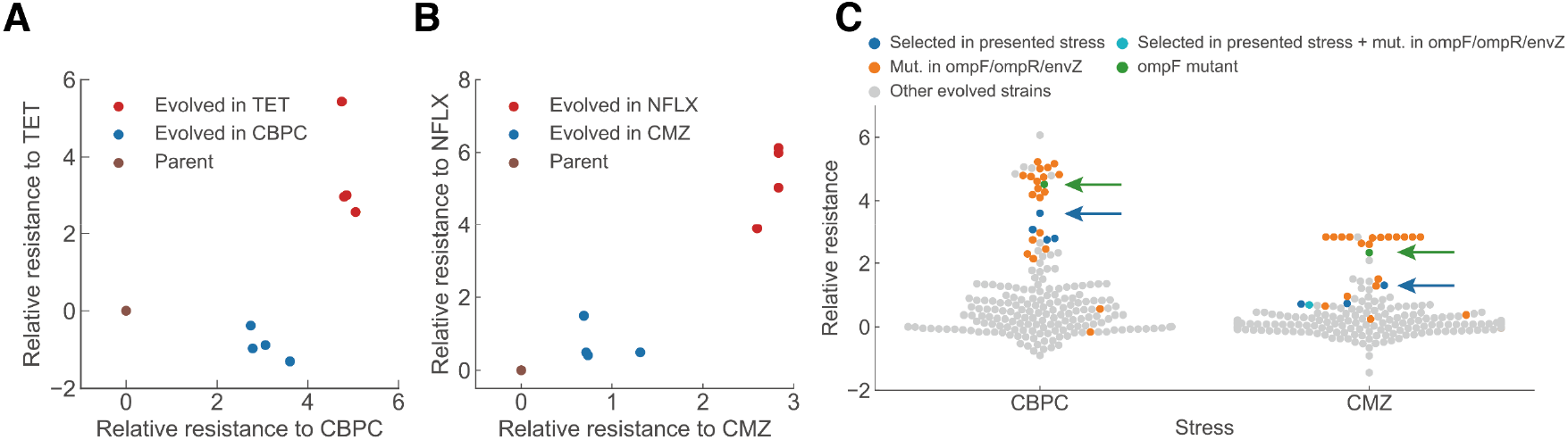
Decelerated evolution against cell wall synthesis inhibitors. (A, B) Decelerated evolution observed within the evolved strains. Relative log_2_ (IC_50_) for evolved strains in CBPC and TET (A), and relative log_2_ (IC_50_) for evolved strains in norfloxacin (NFLX) and cefmetazole (CMZ) (B). (C) Relative IC_50_ values for CBPC and CMZ for all 192 evolved strains. Many of the strains which exhibit resistance higher than the CBPC and CMZ evolved strains had a mutation in *ompF* or its regulators *ompR* and *envZ* (orange). The CBPC and CMZ resistance of the *ompF* introduced strain also exhibited higher resistance (green, green arrow) than the CBPC and CMZ evolved strains (blue, cyan, denoted by a blue arrow).

## DISCUSSION

In this study, we performed laboratory evolution of *E. coli* under various stress conditions and obtained evolved strains under these stresses. Combined with interpretable machine learning, our high-throughput phenotypic analysis led to the identification of modular phenotypic classes both in the gene expression space and the stress resistance space, suggesting tight interactions between changes in gene expression and stress resistance. Furthermore, these classes included strains which evolved in different type of stresses, indicating the existence of evolutionary constraints which do not necessarily depend on the stressor’s mechanism of action.

Our resource includes valuable information on evolutionary constraints for antibiotic resistance, and thus, provides important insights for novel clinical strategies. For instance, we found that various antibiotics with different action mechanisms exhibited collateral sensitivity to metabolic inhibitors including LVAL, B.Cl.Ala, and GAH. We also found that strains which evolved in various stresses showed collateral sensitivity to H_2_O_2_ and protamine sulfate (PS) (15/47 and 14/47 stresses, respectively). Collateral sensitivity to H_2_O_2_ was commonly observed for strains in class 9 and 11 (Fig. S2), which might have been caused by the alteration of global gene expression by mutations in *rssB* (class 9) and *sohA* (class 11) since H_2_O_2_ has many cellular targets (Belenky et al., 2015). On the other hand, collateral sensitivity to PS was commonly observed for strains in class 2, 5, and 11 (Fig. S2), where the enhanced drug efflux by *acrR* mutations (class 5) or the decrease in drug uptake by *ompF* associated mutations (class 2 and 9) might have led to collateral sensitivity to PS, which has been shown to target the cytoplasmic membrane in *Salmonella typhimurium* (Aspedon and Groisman, 1996). These results suggest that the perturbation of metabolic activity, reactive oxygen species (ROS) generation, and alteration of cytoplasmic membrane permeabilization could be possible strategies to suppress antibiotic resistance.

Based on the resequencing analysis of the evolved strains, we clarified the effect of single mutations for resistance acquisition through the introduction of representative mutations to the parent genome. As shown in Fig. 3C, D, the patterns of cross-resistance and collateral sensitivity observed in the reconstructed mutants agreed with that of the evolved strains. These results suggest that the observed evolutionary constraints for resistance in the evolved strains were rooted in the acquired mutations. The analysis of reconstructed mutant strains also provided valuable information on the genetic basis of resistance acquisition. It demonstrated that the pattern of resistance acquisition observed in the evolved strains could be characterized by known resistance-conferring genes, such as *acrAB* and *ompF.* Furthermore, the analysis identified novel genetic mechanisms for the stress resistance, such as CBPC, AZT, and DVAL resistance through the mutation in *sohA* (Fig. 3E), which is related to the TA system. We also discovered the contribution of uncharacterized genes to resistance acquisition, such as the *ycbZ* and *yhjE* mutations causing EM, NMNO, EDTA, NVA, and 5.FOA resistances, and PLM, NVA, and DVAL resistances, respectively.

It should be noted that the analyses presented in this study are not without limitations. First, the identification of distinct classes in phenotypic changes (Fig. 2) was based on gene expression changes, and thus, the analysis missed resistance acquiring mechanisms with small gene expression changes. For example, in the evolved strains under 6.MP, mutations in the *hpt* gene, encoding hypoxanthine phosphoribosyltransferase, were commonly fixed (three out of four evolved strains), suggesting the contribution of the mutation to the resistance phenotype to 6.MP. However, these 6.MP resistant strains exhibited little expression changes compared to the parent, and thus, we could not identify the common phenotypic changes through gene-expression based analysis. Such evolved strains with little expression changes were classified to class 12 (Fig. S3). Including mutations fixed in those strains, a detailed description of the mutations conferring resistance for each stress is given in Table S4.

Next, the introduction of single mutations fixed in the evolved strains was not always enough to explain the resistance changes observed in the laboratory evolution. For example, all evolved strains in class 1 (Fig. 2) had mutations in *mprA*, strongly suggesting the contribution of this mutation to the common phenotypic changes in class 1. However, the reconstructed mutant strain of *mprA* exhibited a similar, but significantly different resistance profile for some stresses such as SXZ (Table S2). The differences between the evolved strains and reconstructed mutant strains might suggest either the contribution of multiple mutations and epistasis among them to the resistance changes, or the contribution of non-genetic adaptation which is difficult to explain by the phenotype-genotype mapping presented in this study. Future studies will likely identify the effect of multiple mutations and non-genetic adaptation on the stress resistance, which will enable a better understanding of phenotypic and genotypic constraints on resistance evolution.

## ACKNOWLEDGMENTS

We thank members of the Furusawa laboratory for fruitful discussions. This work was supported in part by Grant-in-Aid for the Japan Society for the Promotion of Science (JSPS) Fellows [18J21942], Grant-in-Aid for Scientific Research (B) [15KT0085, 17H03622], JSPS KAKENHI [16H06279] (PAGS) and Grant-in-Aid for Scientific Research (S) [15H05746] from JSPS.

## AUTHOR CONTRIBUTIONS

Conceptualization, T.M., J.I., and C.F.; Methodology, T.M., J.I., and C.F.; Software, J. I., K.T., and C.F.; Formal Analysis, J.I. and C.F.; Investigation, T.M., H.K., N.S., T.H., and A.S.; Supervision, C.F; Writing – Original Draft, T.M. and J.I.; Writing – Reviewing and Editing, C.F.; Funding Acquisition, C.F.; Project Administration, C.F.

## SUPPLEMENTAL INFORMATION TITLES AND LEGENDS

**Figure S1. Related to Figure 1. Resistance time series for laboratory evolution** Time series of the highest drug concentration in which the cells could grow (transferred concentration) during the laboratory evolution experiment. The drugs are sorted based on the order of Table S1.

**Figure S2. Related to Figure 2. Combinations of stresses which exhibited cross resistance and collateral sensitivity**

(A) The identified combinations of stresses which exhibited either cross resistance or collateral sensitivity for each of the four strains which evolved in the same environment. (B) The combinations of stresses which exhibited either cross resistance or collateral sensitivity for the strains in each class in the supervised principal component analysis (PCA) space. The combinations were detected by the Mann-Whitney U-test (false discovery rate, FDR < 0.05), and the colors indicate the resistance to the stress relative to the parent strain.

**Figure S3. Related to Figure 2. Phenotypic and genotypic characteristics for all 15 supervised PCA classes**

(A) Dendrogram of the result of hierarchical clustering performed in the 36 dimensional supervised PCA space. (B) Gene expression levels of representative genes for each cluster, relative to the parent strain. The genes were selected from the intersection of the top two gene weights for the linear discriminant analysis (LDA) axis and differentially expressed genes (STAR Methods). (C) Half-maximal inhibitory concentrations (IC_50_) values relative to the parent strain. (D) Commonly mutated genes within the evolved strains. Mutated genes enriched for each cluster clarified by Fisher’s exact test (p < 0.01) are presented. Mutated genes that were identified in more than seven strains are also presented. (E) Feature importance of the genes computed by the random forest regression model. (F) Class dissimilarity in the IC_50_ space for the 15 classes which were defined by hierarchical clustering based on IC_50_ values, supervised PCA expression space, mutations, and full gene expression space, respectively. The mean and standard deviation for the class dissimilarity for 10 runs of randomly clustered results are also shown.

**Figure S4. Related to Figure 2 and Figure 3. Combinations of stresses which exhibited cross resistance, collateral sensitivity**

(A) Stresses which exhibited either resistance or sensitivity for the 64 reconstructed mutant strains (Mann-Whitney U-test, p < 0.05). Colors indicate the stress resistance relative to the parent strain. (B) and (C) Stress resistance relative to the parent strain for strains in class 9 and class 11, respectively. Resistance levels for the *rssB* and *sohA* mutant are also shown for comparison. The p-values were calculated by the Mann-Whitney U-test, which compared the evolved strains in the corresponding class and the parent strains.

**Figure S5. Related to Figure 4. Neither fitness trade-offs nor epistasis explains decelerated evolution**

(A) and (B) Growth rates for the parent strain, *ompF* mutated strain, and the *sohA* mutant. Growth rates were measured in 24 concentration levels of cefmetazole (CMZ) and carbenicillin (CBPC), respectively. (C) IC_50_ levels measured for different stresses for the parent strain, *ompF* mutated strain, *sohA* mutated strain, and the *sohA*/*ompF* doubled mutated strain, respectively.

**Table S1. Related to Figure 1: List of the chemicals used in this study**

**Table S2. Related to Figure 2: All relative half-maximal inhibitory concentrations (IC_50_s) for the 47 stresses of each evolved strain and reconstructed mutant strain**

**Table S3. Related to Figure 2: Transcriptome data of evolved strains.**

**Table S4. Related to Figure 2: All identified mutations in the evolved strains**

All identified mutations in the evolved strains are shown in sheet 1. Representative genes in which non-synonymous mutations or ins/dels were commonly fixed in the evolved strains are listed in sheet 2. Primers used for the constructions of the reconstructed mutant strains are also shown in sheet 2. Detailed descriptions of common mutations identified in the evolved strains are shown in sheet 3.

**Table S5. Related to Figure 3: Cross-resistances and collateral sensitivities observed in the 64 reconstructed mutant strains**

Chemicals that were identified as significantly increased or decreased IC_50_ values (Mann-Whitney U-test, false discovery rate (FDR) < 5%) in the all reconstructed mutant strains are shown.

## STAR METHODS

### Lead Contact and Materials Availability

Further information and requests for resources and regents should be directed to and will be fulfilled by the Lead Contact, Chikara Furusawa. All unique/stable reagents generated in this study are available from the Lead Contact without restriction.

### Experimental Model and Subject Details

#### Bacterial strains and growth media

The insertion sequence (IS)-free *E. coli* strain MDS42 (Pósfai et al., 2006) was purchased from Scarab Genomics (Scarab Genomics, Madison, Wisconsin, USA). *E. coli* cells were cultured in modified M9 minimal medium containing 17.1 g/L Na_2_HPO_4_·12H_2_O, 3.0 g/L KH_2_PO_4_, 5.0 g/L NaCl, 2.0 g/L NH_4_Cl, 5.0 g/L glucose, 14.7 mg/L CaCl_2_·2H_2_O, 123.0 mg/L MgSO_4_·7H_2_O, 2.8 mg/L FeSO_4_·7H_2_O, and 10.0 mg/L thiamine hydrochloride (pH 7.0) (Mori et al., 2011) plus 15 μg/mL erythromycin. To construct mutant strains, LB medium and Terrific broth (TB) were used. LB medium contained 10 g/L Bacto tryptone, 5 g/L Bacto yeast extract, and 5 g/L NaCl. TB contained 12 g/L Bacto tryptone, 24 g/L Bacto yeast extract, 4 g/L glycerol, 2.32 g/L KH_2_PO_4_, and 12.54 g/L K_2_HPO_4_.

#### Laboratory evolution

Supplementary Table S1 lists all of the chemicals that were used in this study and the solvents in which they were dissolved to prepare stock solutions. Chemicals that did not dissolve in the modified M9 medium were added to it at a > 20-fold dilution. Cell cultivation, optical density (OD) measurements, and serial dilutions were performed for each chemical using an automated culture system (Maeda et al., 2019) consisting of a Biomek^®^ NX span-8 laboratory automation workstation (Beckman Coulter, Brea, California, USA) in a clean booth connected to a microplate reader (FilterMax F5; Molecular Devices, San Jose, California, USA), a shaker incubator (STX44; Liconic, Mauren, Liechtenstein), and a microplate hotel (LPX220, Liconic). A movie of the automated culture system is available on YouTube (https://www.youtube.com/watch?v=4k6qCN7ppsk).

Before laboratory evolution, the MDS42 strain was cultivated in modified M9 medium without chemicals used for laboratory evolution for 96 h (approximately 90 generations) to acclimatize them to the modified M9 medium. After the 96-h cultivation, a glycerol stock of the cells was stored as the parent strain at –80 °C for further experiments. Six independent culture lines were evolved in parallel for each chemical. 384-well microplates containing 45 μL modified M9 medium per well and a 2^0.25^-fold chemical gradient in 22 dilutions, were used. To start laboratory evolution, MDS42 cells were firstly inoculated from the frozen glycerol stock into the modified M9 medium and cultivated 24 h at 34 °C with shaking. The OD_620_ values of the precultures were measured using the automated culture system, and precultured cells that were calculated to have initial OD_620_ values of 0.0003 were inoculated into each well (5 μL of diluted overnight culture into 45 μL of medium per well) of the 384-well microplates and cultivated with agitation at 300 rotations/min at 34 °C. Every 24 h of cultivation, cell growth was monitored by measuring the OD_620_ of each well. We defined a well whose OD_620_ was > 0.09 as a well in which cells grew. The automated culture system automatically selected the defined well with the highest chemical concentration in which cells could grow; the cells in the selected well were diluted to an OD_620_ of 0.0003 and transferred to new plates containing fresh medium and chemical gradients. During and after the laboratory evolution cells were stored as glycerol stocks at −80 °C. We isolated a single clone on a modified M9 agar plate without chemicals to use for laboratory evolution from the endpoint culture and confirmed that the IC_50_ of the isolated clone was almost identical to that of the corresponding population in the endpoint culture. We used the isolated single clones as the evolved strains for further analysis.

### Method Details

#### Total RNA purification

Total cellular RNA was isolated as follows. Cells were inoculated from the frozen glycerol stock into 96-well microplates containing 200 μL of modified M9 medium without chemicals and cultivated 24 h at 34 °C with agitation at 900 rotations/min. The precultures were diluted to an OD_620_ of 3.3 × 10^−5^ - 4.0 × 10^−4^ into 200 μL of fresh modified M9 medium in 96-well microplates and cultured. Cell growth was monitored by measuring the OD_600_ of each well using the 1420 ARVO microplate reader (PerkinElmer Inc.). After 12 to 21 h incubation, the exponentially growing cultures (OD_600_ in the 0.072-0.135 range) were selected and treated immediately by adding an equal volume of ice-cold ethanol containing 10% (w/v) phenol to stabilize the cellular RNA. The cells were harvested by centrifugation at 20,000 × *g* at 4 °C for 5 min, and the pelleted cells were stored at –80 °C before RNA extraction. Total cellular RNA was isolated using the RNeasy Mini Kit (Qiagen) according to the manufacturer’s instructions. Total RNA was treated with DNase I at room temperature for 15 min and the purified RNA samples were stored at –80 °C until the microarray experiments were performed.

#### Transcriptome analysis using microarrays

Microarray experiments were performed as described previously (Suzuki et al., 2014) using a custom-designed Agilent 8 × 60 K array for *E. coli* W3110 that included 12 probes for each gene. Briefly, 100 ng of each purified total RNA sample was labeled using the Low Input Quick Amp WT Labeling Kit (Agilent Technologies, Santa Clara, California, USA) with Cyanine3 (Cy3) according to the manufacturer’s instructions. Cy3-labeled cRNAs were fragmented and hybridized to the microarray for 17 h at 65 °C in a hybridization oven (Agilent Technologies); the microarray was then washed and scanned according to the manufacturer’s instructions. Microarray image analysis was performed using Feature Extraction version 10.7.3.1 (Agilent Technologies) and expression levels were normalized using the quantile normalization method (Bolstad et al., 2003). Each experiment was performed in triplicate, starting from independent cultures. The microarray data have been submitted to the National Center for Biotechnology Information’s Gene Expression Omnibus functional genomics data repository under accession number GSE137348.

#### qRT-PCR

The mRNA was quantified using the CFX96 Real-Time PCR Detection System (Bio-Rad). qRT-PCR analysis was performed using the following primer sets: 5’-CGACCTGTTAGACGCTGATT-3’ and 5’-GTTCAGCGGTAACACGGATT-3’ (*gapA*), 5’-CGCGCTTATCGTGAAGAGGC-3’ and 5’-GTGCCGCTGTCGGTCAGTAA-3’ (*tnaA*), 5’-GGCAATGGCGACATGACCTA-3’ and 5’-GCGCCTTCAGAGTTGTTACC-3’ (*ompF*), 5’-GGCAAGCACCATCAGACTTA-3’ and 5’-CCAGACCTGAGCTGCGTTGT-3’ (*sodB*). A 50 ng total RNA sample was used for each RT-PCR with each primer pair using the iTaq Universal SYBR Green One-Step Kit (Bio-Rad) according to the manufacturer’s instructions. The target gene transcripts were normalized to the reference gene transcript (*gapA*) from the same RNA sample. Each gene was analyzed using RNA isolated from three independent samples. The cycle threshold (CT) for each sample was generated according to the procedures described in the CFX96 Real-Time PCR Detection System user guide.

#### Preparation of genomic DNA

Stock strains were inoculated in 5 mL of modified M9 medium without chemicals in test tubes for 24 h at 34 °C and 150 rpm using water bath shakers (Personal-11, Taitec Co.). 300 μg/mL rifampicin (RFP) was subsequently added and the culture was continued for a further 3 h to inhibit the initiation of DNA replication. The cells were collected by centrifugation at 25 °C and 20,000 × *g* for 5 min and the pelleted cells were stored at – 80 °C before genomic DNA purification. Genomic DNA was isolated and purified using a DNeasy Blood & Tissue Kit (Qiagen) in accordance with the manufacturer’s instructions. The quantity and purity of the genomic DNA were determined by measuring the absorbance at 260 nm and calculating the ratio of absorbance at 260 and 280 nm (A_260_/_280_) using a NanoDrop ND-2000 spectrophotometer, respectively. The A_260_/_280_ values of all the samples were confirmed to be greater than 1.7. The purified genomic DNAs were stored at −30 °C before use.

#### Genome sequence analysis using Illumina HiSeq system

Genome sequence analyses using the Illumina HiSeq System were described previously (Horinouchi et al., 2017). A 150-bp paired-end library was generated according to the Illumina protocol and sequenced using Illumina HiSeq. In this study, 192 samples with different barcodes were mixed and then sequenced, which resulted in approximately 140-fold coverage on average. The quality of the sequence data was first assessed using FastX-Toolkit 0.0.13.2 (http://hannonlab.cshl.edu/fastx_toolkit), and the raw reads were trimmed using PRINSEQ (Schmieder and Edwards, 2011), whereby both ends with quality scores lower than Q20 were trimmed. The potential nucleotide differences were validated using BRESEQ (Deatherage and Barrick, 2014).

#### Introduction of identified mutations into the parent strain

To construct mutant strains, the identified mutations were introduced into the parental strain by pORTMAGE (Nyerges et al., 2016). pORTMAGE-4 was a gift from Dr. Csaba Pál (Addgene plasmid # 72679; http://n2t.net/addgene:72679; RRID:Addgene_72679). The introduced mutations and the DNA oligonucleotides used in this study are listed in Table S4. Briefly, MDS42 strain harboring pORTMAGE-4 plasmid was inoculated into 5 mL LB media in the presence of 20 μg/mL chloramphenicol and incubated at 30 °C and 150 rpm. Overnight cultures were diluted 1:100 in chloramphenicol supplemented LB media and grown at 30 °C with agitation at 1,000 rotations/min. After 1.5 h incubation, the exponentially growing cultures (OD_600_ = 0.4-0.6) were further incubated at 42 °C for 5 min at 1,000 rotations/min to induce λ Red protein expression. Cells were then immediately chilled on ice for 10 min. Electrocompetent cells were prepared by washing and pelleting twice in ice-cold sterile MilliQ water. Each MAGE oligo (90-mer oligonucleotide containing desired mutation and four phosphorothioated bases at the 5’ termini) at 2.5 μM final concentration was introduced by electroporation. For gene inactivation, except for *acrR*, NheI site containing the TAG stop codon was introduced immediately downstream of its start codon and one base was inserted to introduce a frameshift mutation. To construct the *acrR* mutant strain, a NheI site was introduced at 24 bp downstream of the start codon. After electroporation, 1 mL TB medium was added, and cells were incubated at 30 °C for 1 h. At this point, cells were subjected to additional MAGE cycles. After the fourth MAGE cycle, cells were diluted 1:1000 and plated onto LB media. After 18 h incubation at 30 °C, single colonies were picked, and corresponding genomic regions were amplified by PCR and then verified by Sanger sequencing or NheI digestion to select the desired mutant strains. The selected mutants were further cultivated in TB medium to eliminate the pORTMAGE-4 plasmid. The correctness of the constructed mutants was further confirmed by Sanger sequencing.

#### Prediction and gene selection using a random forest model

A random forest model was constructed to predict the relative IC_50_s for each of the 47 stresses from the 4,492 gene expression levels for all 192 evolved strains. Here, the scikit-learn implementation of the random forest regressor was used. The log_10_-transformed gene expression levels were used for prediction. Since the relative IC_50_ values had different scales, we normalized the data between stresses by multiplying 1, 0.5, or 0.25 depending on the maximum fold changes of the IC_50_s for the evolved strains against the corresponding chemicals. This normalization corresponds to the chemical gradient dilution steps used for the experimental determination of the IC_50_s. To avoid overfitting, we applied a grid search over the number of trees (16 values between 10 and 40) and the max depth of each tree (60 values between 20 and 1,200), using a 4-fold cross validation method. The set of hyperparameters which provided the lowest mean squared prediction error, (number of trees, max depth) = (300, 18), was selected for further analysis. Using the optimal hyperparameters, we trained the random forest on the whole dataset in order to extract the ranking of relative gene importance for IC_50_ prediction. The relative gene importance was evaluated by the decrease of the mean squared prediction error at each branch of the tree through the feature_importance attribute of the RandomForestRegressor function. The 213 genes which had high feature importance deviating from the trend of exponential decay were selected for supervised PCA (Fig. S3).

#### Supervised PCA and hierarchical clustering

The expression profiles of the 213 genes, selected through the random forest model, were used for PCA (supervised PCA). Hierarchical clustering was applied to the supervised PCA space using Ward’s method. The number of classes was determined by the elbow method using class dissimilarity (*W*_*n*_) as a criterion. *W*_*n*_ was defined as 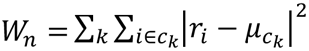, where *k*, *c*_*k*_, *μc*_*k*_ is the class’s index, the set of elements for each class, and class k’s centroid, respectively, and *r*_*i*_,*i* represents the location of each strain, and its index, respectively. For each number of classes *n*, we calculated *W*_*n*_, and its derivative (*W*_*n*_ − *W*_*n*-1_) in the resistance space and searched for the number of classes where the derivative of *W*_*n*_ sharply decreased. As a result, the optimal number of classes was determined to be 15. The results of hierarchical clustering including all 15 classes are given in Fig. S3.

#### LDA

To investigate the representative gene expression levels for each class, we applied LDA to the transcriptome data. LDA was performed by using the LinearDiscriminantAnalysis function from the scikit-learn package. The strains were given binary labels for LDA: one for the strains which belonged to the class of interest, and zero for the other strains. To extract the important genes which characterized each class, we looked for the top weighted genes in each LDA axis, which corresponded to the genes which contributed to the decision boundary for the binary labeled strains. We further selected the genes which had more than a two-fold change in gene expression compared with the parent strain, within the top weighted genes in the LDA axis.

#### Comparing class dissimilarities

To evaluate how accurately the neighboring relationship in the resistance space was conserved in the supervised PCA space, we calculated *W*_*n*_ in the resistance space based on the clustering results in the resistance space, supervised PCA space, mutation space, and the full expression space, respectively. For the resistance space, hierarchical clustering was applied based on the 47 relative IC_50_s, to cluster the 192 strains to 15 classes. For the mutation space, hierarchical clustering was applied based on the one-hot encoding which reflects the information of the presence of a mutation. For the expression space, hierarchical clustering was applied to the whole 4,492-dimension gene expression space. To construct a baseline, the class dissimilarity was calculated for randomly clustered classes in the resistance space as well. The results are given in Fig. S3.

#### Quantification and statistical analysis

To determine IC_50_s, serial dilutions of each chemical were prepared in 384-well microplates using the modified M9 medium with 2^0.25^-, 2^0.5^-, or 2-fold chemical gradients in 22 dilution steps. The chemical gradients depended on the maximum fold changes of the IC_50_s for the evolved strains against the correspondent chemicals. Culture conditions for IC_50_ determination was the same as for laboratory evolution. After 24 h cultivation in the 384-well microplates containing serially diluted chemicals, the OD_620_ of the cultures was measured. To obtain the IC_50_ values, the OD_620_ values for the dose-response series were fitted to the following sigmoidal model: *f*(*x*) = *a*/[1 + exp *b*(log_2_ *x* − log_2_ *IC*_*50*_)] + *c*, where *x* and *f(x)* represent the concentration of antibiotics and observed OD_620_ values, respectively, and a, b, c, and IC_50_ are fitting parameters. The fitting was performed using optimize.curve_fit in the SciPy package (Virtanen et al., 2019). The relative IC_50_s were computed by comparing the IC_50_ of each evolved strain to the mean of 13 independent replicas of the parent strain.

Cross resistance and collateral sensitivities of stresses were investigated through the Mann-Whitney U-test. To detect collateral relationships between stress A and B, we compared the IC_50_ values of stress B of the four strains that evolved in stress A to that of the parent strain (13 independent replicas). The p-values were obtained using “wilcox.test” (correct=F, exact=T) in the stats package of R software. We further applied the Benjamini-Hochberg FDR control to these p-values so that FDR < 5%.

#### Data and code availability

The program for supervised PCA and hierarchical clustering is available at https://github.com/jiwasawa/evolved-strain-analysis/. All microarray data are available in the National Center for Biotechnology Information’s Gene Expression Omnibus functional genomics data repository under accession number GSE137348. The raw sequence data of genome sequence analyses are available in the DDBJ Sequence Read Archive under the accession number DRA006396.

## REFERENCES

Aspedon, A., and Groisman, E.A. (1996). The antibacterial action of protamine: evidence for disruption of cytoplasmic membrane energ ization in *Salmonella typhimurium*. Microbiology 142, 3389–3397.

Balagué, C., and Véscovi, E.G. (2001). Activation of multiple antibiotic resistance in uropathogenic *Escherichia coli* strains by aryloxoalcanoic acid compounds. Antimicrob. Agents Chemother. 45, 1815–1822.

Barbosa, C., Trebosc, V., Kemmer, C., Rosenstiel, P., Beardmore, R., Schulenburg, H., and Jansen, G. (2017). Alternative evolutionary paths to bacterial antibiotic resistance cause distinct collateral effects. Mol. Biol. Evol. 34, 2229–2244.

Belenky, P., Ye, J.D., Porter, C.B.M., Cohen, N.R., Lobritz, M.A., Ferrante, T., Jain, S., Korry, B.J., Schwarz, E.G., Walker, G.C., et al. (2015). Bactericidal antibiotics induce toxic metabolic perturbations that lead to cellular damage. Cell Rep. 13, 968–980.

Bolstad, B.M., Irizarry, R.A., Astrand, M., and Speed, T.P. (2003). A comparison of normalization methods for high density oligonucleotide array data based on variance and bias. Bioinformatics 19, 185–193.

Chevereau, G., Dravecká, M., Batur, T., Guvenek, A., Ayhan, D.H., Toprak, E., and Bollenbach, T. (2015). Quantifying the determinants of evolutionary dynamics leading to drug resistance. PLoS Biol. 13, e1002299.

Conrad, T.M., Lewis, N.E., and Palsson, B.Ø. (2011). Microbial laboratory evolution in the era of genome-scale science. Mol. Syst. Biol. 7, 509.

Deatherage, D.E., and Barrick, J.E. (2014). Identification of mutations in laboratory-evolved microbes from next-generation sequencing data using *breseq*. In: Sun L., Shou W. (eds) Engineering and Analyzing Multicellular Systems. Methods in Molecular Biology (Methods and Protocols), vol 1151. Humana Press, New York, NY.

Delcour, A.H. (2009). Outer membrane permeability and antibiotic resistance. Biochim. Biophys. Acta 1794, 808–816.

Ferenci, T. (2005). Maintaining a healthy SPANC balance through regulatory and mutational adaptation. Mol. Microbiol. 57, 1–8.

Furusawa, C., Horinouchi, T., and Maeda, T. (2018). Toward prediction and control of antibiotic-resistance evolution. Curr. Opin. Biotechnol. 54, 45–49.

Gagarinova, A., Stewart, G., Samanfar, B., Phanse, S., White, C.A., Aoki, H., Deineko, V., Beloglazova, N., Yakunin, A.F., Golshani, A., et al. (2016). Systematic genetic screens reveal the dynamic global functional organization of the bacterial translation machinery. Cell Rep. 17, 904–916.

Gibson, K.E., and Silhavy, T.J. (1999). The LysR homolog LrhA promotes RpoS degradation by modulating activity of the response regulator SprE. J Bacteriol 181, 563–571.

Girgis, H.S., Hottes, A.K., and Tavazoie, S. (2009). Genetic architecture of intrinsic antibiotic susceptibility. PLoS One 4, e5629.

Harder, K.J., Nikaido, H., and Matsuhashi, M. (1981). Mutants of *Escherichia coli* that are resistant to certain beta-lactam compounds lack the *ompF* porin. Antimicrob. Agents Chemother. 20, 549–552.

Horinouchi, T., Minamoto, T., Suzuki, S., Shimizu, H., and Furusawa, C. (2014). Development of an Automated Culture System for Laboratory Evolution. J. Lab. Autom. 19, 478–482.

Horinouchi, T., Suzuki, S., Kotani, H., Tanabe, K., Sakata, N., Shimizu, H., and Furusawa, C. (2017). Prediction of cross-resistance and collateral sensitivity by gene expression profiles and genomic mutations. Sci. Rep. 7, 14009.

Imamovic, L., Ellabaan, M.M.H., Dantas Machado, A.M., Citterio, L., Wulff, T., Molin, S., Krogh Johansen, H., and Sommer, M.O.A. (2018). Drug-driven phenotypic convergence supports rational treatment strategies of chronic infections. Cell 172, 121–134.

Imlay, J.A., and Linn, S. (1987). Mutagenesis and stress responses induced in *Escherichia coli* by hydrogen peroxide. J. Bacteriol. 169, 2967–2976.

Jiao, Y.J., Baym, M., Adrian, V., and Kishony, R. (2016). Population diversity jeopardizes the efficacy of antibiotic cycling. BioRxiv.

Lässig, M., Mustonen, V., and Walczak, A.M. (2017). Predicting evolution. Nat. Ecol. Evol. 1, 77.

Lázár, V., Nagy, I., Spohn, R., Csörgő, B., Györkei, Á., Nyerges, Á., Horváth, B., Vörös, A., Busa-Fekete, R., Hrtyan, M., et al. (2014). Genome-wide analysis captures the determinants of the antibiotic cross-resistance interaction network. Nat. Commun. 5, 4352.

Lee, H.H., Molla, M.N., Cantor, C.R., and Collins, J.J. (2010). Bacterial charity work leads to population-wide resistance. Nature 467, 82–85.

Li, X.-Z., and Nikaido, H. (2009). Efflux-mediated drug resistance in bacteria: an update. Drugs 69, 1555–1623.

Lomovskaya, O., Lewis, K., and Matin, A. (1995). EmrR is a negative regulator of the *Escherichia coli* multidrug resistance pump EmrAB. J. Bacteriol. 177, 2328–2334.

Ma, D., Cook, D.N., Alberti, M., Pon, N.G., Nikaido, H., and Hearst, J.E. (1993). Molecular Cloning and Characterization of *acrA* and *acrE* Genes of *Escherichia coli*. J. Bacteriol. 175, 6299–6313.

Maeda, T., Horinouchi, T., Sakata, N., Sakai, A., and Furusawa, C. (2019). High-throughput identification of the sensitivities of an *Escherichia coli* Δ*recA* mutant strain to various chemical compounds. J. Antibiot. (Tokyo). 72, 566–573.

May, M. (2014). Drug development: Time for teamwork. Nature *509*, S4–S5.

Mazzariol, A., Cornaglia, G., and Nikaido, H. (2000). Contributions of the AmpC beta-lactamase and the AcrAB multidrug efflux system in intrinsic resistance of *Escherichia coli* K-12 to beta-lactams. Antimicrob. Agents Chemother. 44, 1387–1390.

Mori, E., Furusawa, C., Kajihata, S., Shirai, T., and Shimizu, H. (2011). Evaluating 13C enrichment data of free amino acids for precise metabolic flux analysis. Biotechnol. J. 6, 1377–1387.

Munck, C., Gumpert, H.K., Wallin, A.I.N., Wang, H.H., and Sommer, M.O.A. (2014). Prediction of resistance development against drug combinations by collateral responses to component drugs. Sci. Transl. Med. 6, 262ra156.

Nichol, D., Rutter, J., Bryant, C., Hujer, A.M., Lek, S., Adams, M.D., Jeavons, P., Anderson, A.R.A., Bonomo, R.A., and Scott, J.G. (2019). Antibiotic collateral sensitivity is contingent on the repeatability of evolution. Nat. Commun. 10, 334.

Norioka, S., Ramakrishnan, G., Ikenaka, K., and Inouye, M. (1986). Interaction of a Transcriptional Activator, OmpR, with Reciprocally Osmoregulated Genes,. J. Biol. Chem. 261, 17113–17119.

Nyerges, Á., Csörgő, B., Nagy, I., Bálint, B., Bihari, P., Lázár, V., Apjok, G., Umenhoffer, K., Bogos, B., Pósfai, G., et al. (2016). A highly precise and portable genome engineering method allows comparison of mutational effects across bacterial species. Proc. Natl. Acad. Sci. 113, 2502–2507.

O’Neill, J. (2016). Tackling drug-resistant infections globally: Final report and recommendations. https://amr-review.org/sites/default/files/160518_Final%20paper_with%20cover.pdf.

Okusu, H., Ma, D., and Nikaido, H. (1996). AcrAB efflux pump plays a major role in the antibiotic resistance phenotype of *Escherichia coli* multiple-antibiotic-resistance (Mar) mutants. J. Bacteriol. 178, 306–308.

Palmer, A.C., and Kishony, R. (2013). Understanding, predicting and manipulating the genotypic evolution of antibiotic resistance. Nat. Rev. Genet. 14, 243–248.

Palmer, A.C., and Kishony, R. (2014). Opposing effects of target overexpression reveal drug mechanisms. Nat. Commun. 5, 4296.

Pósfai, G., Plunkett III, G., Fehér, T., Frisch, D., Keil, G.M., Umenhoffer, K., Kolisnychenko, V., Stahl, B., Sharma, S.S., de Arruda, M., et al. (2006). Emergent properties of reduced-genome *Escherichia coli*. Science. 312, 1044–1046.

Radzikowski, J.L., Vedelaar, S., Siegel, D., Ortega, Á.D., Schmidt, A., and Heinemann, M. (2016). Bacterial persistence is an active σS stress response to metabolic flux limitation. Mol. Syst. Biol. 12, 882.

Schmidt, O., Schuenemann, V.J., Hand, N.J., Silhavy, T.J., Martin, J., Lupas, A.N., and Djuranovic, S. (2007). *prlF* and *yhaV* encode a new toxin-antitoxin system in *Escherichia coli*. J. Mol. Biol. 372, 894–905.

Schmieder, R., and Edwards, R. (2011). Quality control and preprocessing of metagenomic datasets. Bioinformatics 27, 863–864.

Shibai, A., Takahashi, Y., Ishizawa, Y., Motooka, D., Nakamura, S., Ying, B.-W., and Tsuru, S. (2017). Mutation accumulation under UV radiation in *Escherichia coli*. Sci. Rep. 7, 14531.

Suk, J.E., Vaughan, E.C., Cook, R.G., and Semenza, J.C. (2019). Natural disasters and infectious disease in Europe : a literature review to identify cascading risk pathways. Eur. J. Public Health 1–8.

Suzuki, S., Horinouchi, T., and Furusawa, C. (2014). Prediction of antibiotic resistance by gene expression profiles. Nat. Commun. 5, 5792.

Tenaillon, O., Tenaillon, O., Rodríguez-verdugo, A., Gaut, R.L., Mcdonald, P., Bennett, A.F., Long, A.D., and Gaut, B.S. (2012). The molecular diversity of adaptive convergence. Science. 335, 457–461.

Toprak, E., Veres, A., Michel, J.-B., Chait, R., Hartl, D.L., and Kishony, R. (2011). Evolutionary paths to antibiotic resistance under dynamically sustained drug selection. Nat. Genet. 44, 101–105.

Virtanen, P., Gommers, R., Oliphant, T.E., Haberland, M., Reddy, T., Cournapeau, D., Burovski, E., Peterson, P., Weckesser, W., Bright, J., et al. (2019). SciPy 1. 0 — Fundamental algorithms for scientific computing in Python. ArXiv:1907.10121.

Yelin, I., and Kishony, R. (2018). Antibiotic Resistance. Cell *172*, 1136–1136.e1.

Yoshida, M., Reyes, S.G., Tsuda, S., Horinouchi, T., Furusawa, C., and Cronin, L. (2017). Time-programmable drug dosing allows the manipulation, suppression and reversal of antibiotic drug resistance in vitro. Nat. Commun. 8, 15589.

Zampieri, M., Enke, T., Chubukov, V., Ricci, V., Piddock, L., and Sauer, U. (2017). Metabolic constraints on the evolution of antibiotic resistance. Mol. Syst. Biol. 13, 917.

Zhen, X., Lundborg, C.S., Sun, X., Hu, X., and Dong, H. (2019). Economic burden of antibiotic resistance in ESKAPE organisms: a systematic review. Antimicrob. Resist. Infect. Control 8, 137.

