## Supplemental Figures for "High-throughput laboratory evolution and evolutionary constraints in *Escherichia coli*"

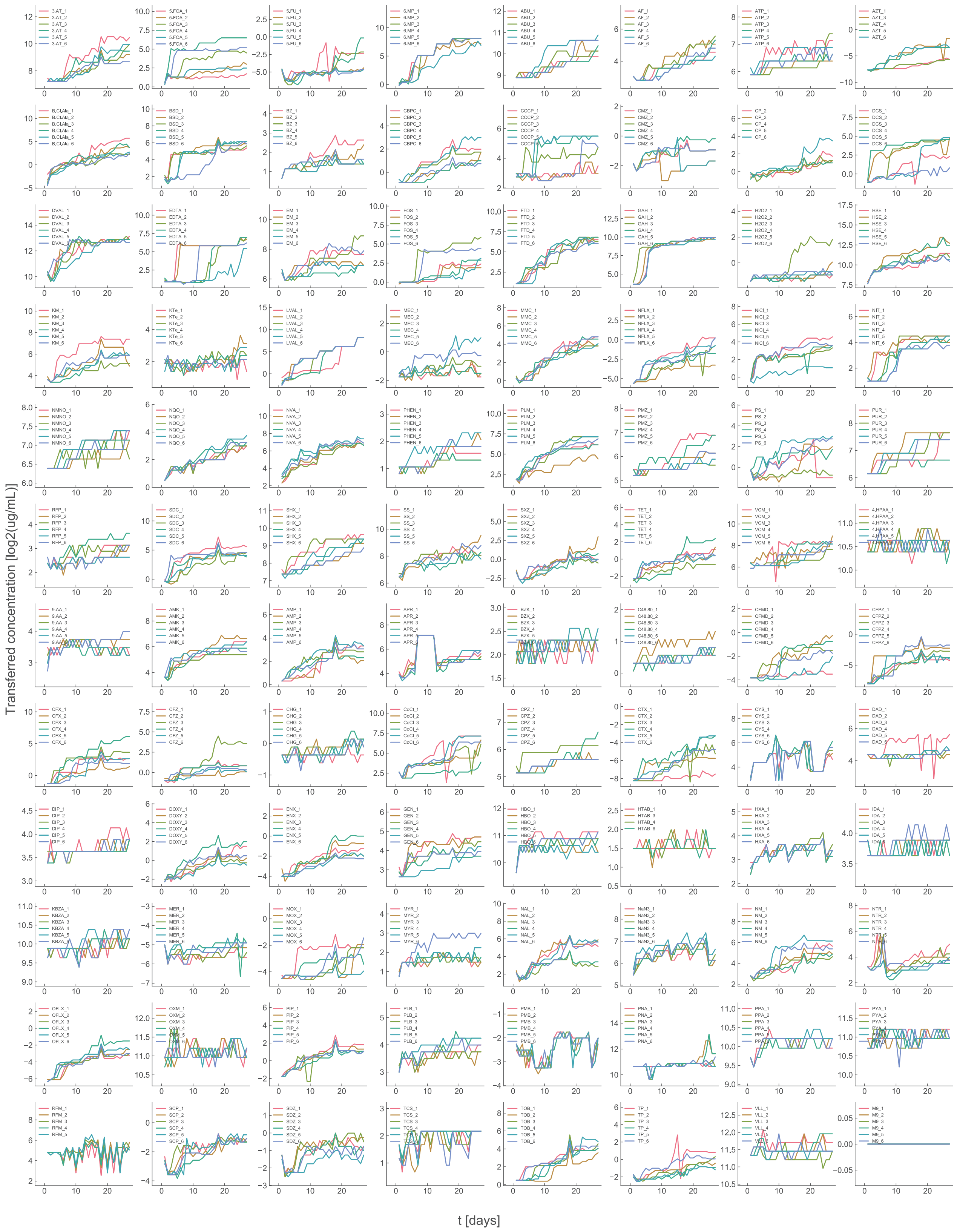

**Figure S1. Related to Figure 1. Resistance time series for laboratory evolution**

Time series of the highest drug concentration in which the cells could grow (transferred concentration) during the laboratory evolution experiment. The drugs are sorted based on the order of Table S1.

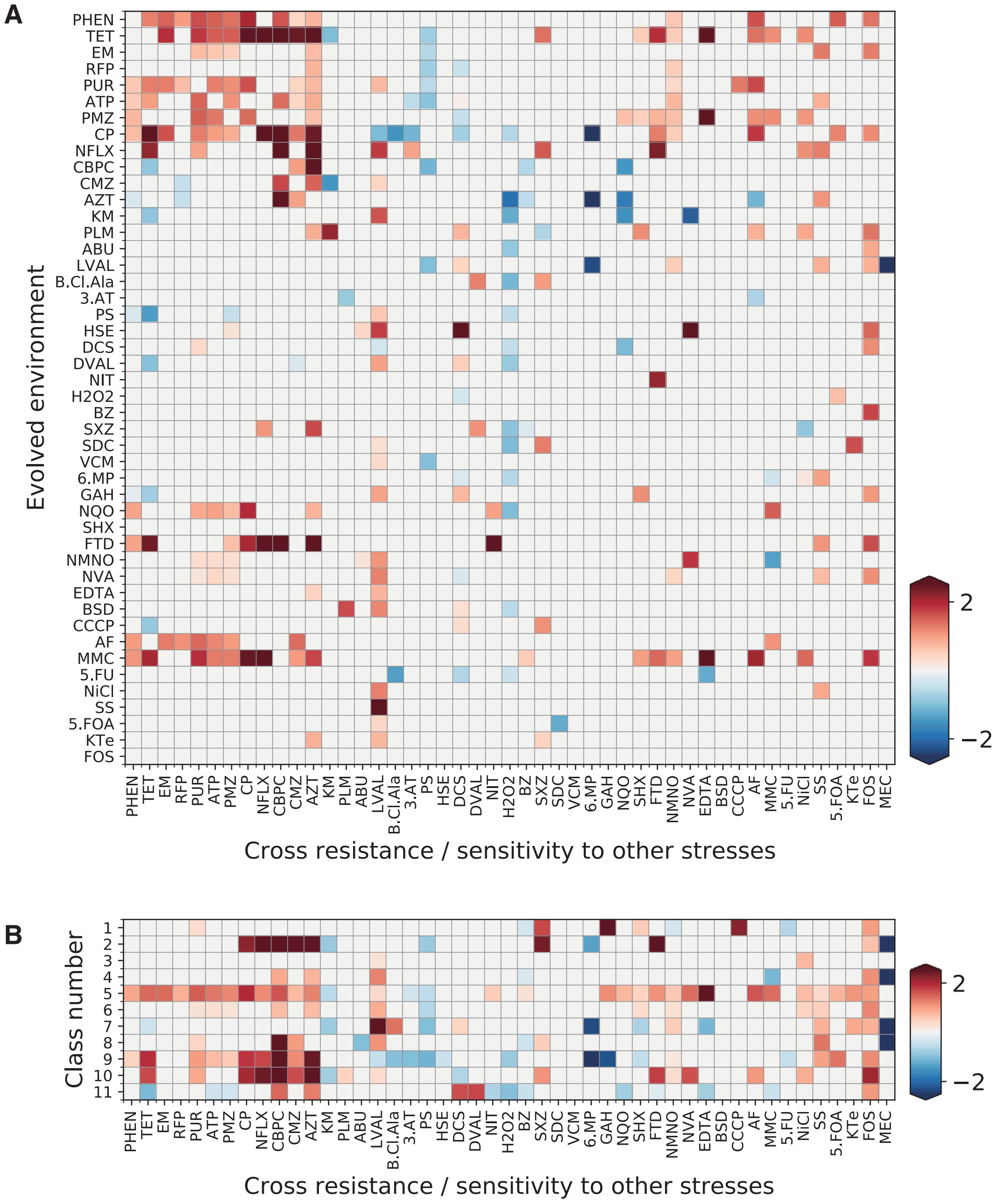


**Figure S2. Related to Figure 2. Combinations of stresses which exhibited cross resistance and collateral sensitivity**

(A) The identified combinations of stresses which exhibited either cross resistance or collateral sensitivity for each of the four strains which evolved in the same environment. (B) The combinations of stresses which exhibited either cross resistance or collateral sensitivity for the strains in each class in the supervised principal component analysis (PCA) space. The combinations were detected by the Mann-Whitney U-test (false discovery rate, FDR < 0.05), and the colors indicate the resistance to the stress relative to the parent strain.

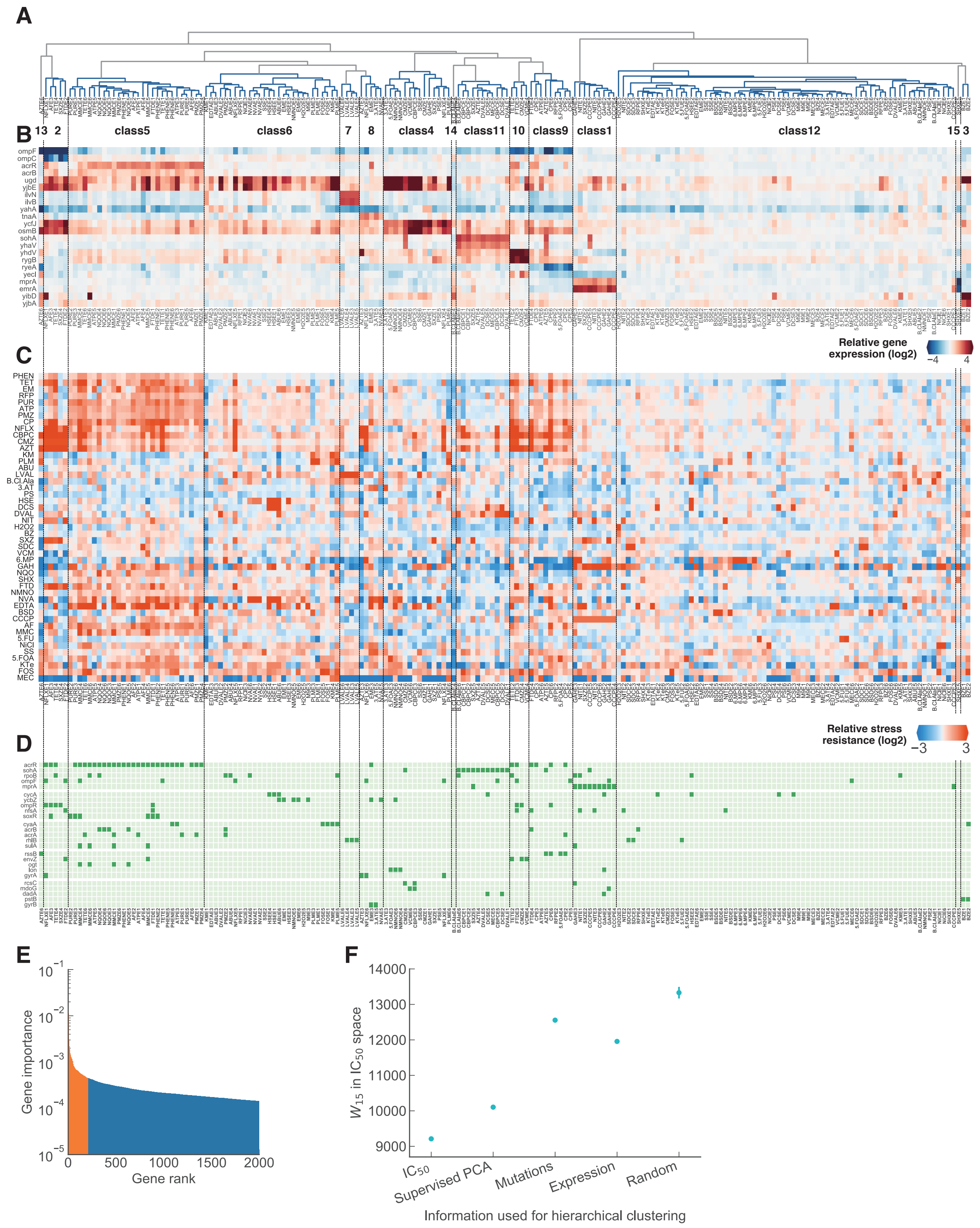


**Figure S3. Related to Figure 2. Phenotypic and genotypic characteristics for all 15 supervised PCA classes**

(A) Dendrogram of the result of hierarchical clustering performed in the 36 dimensional supervised PCA space. (B) Gene expression levels of representative genes for each cluster, relative to the parent strain. The genes were selected from the intersection of the top two gene weights for the linear discriminant analysis (LDA) axis and differentially expressed genes (STAR Methods). (C) Half-maximal inhibitory concentrations (IC50) values relative to the parent strain. (D) Commonly mutated genes within the evolved strains. Mutated genes enriched for each cluster clarified by Fisher's exact test ($p < 0.01$) are presented. Mutated genes that were identified in more than seven strains are also presented. (E) Feature importance of the genes computed by the random forest regression model. (F) Class dissimilarity in the IC50 space for the 15 classes which were defined by hierarchical clustering based on IC50 values, supervised PCA expression space, mutations, and full gene expression space, respectively. The mean and standard deviation for the class dissimilarity for 10 runs of randomly clustered results are also shown.

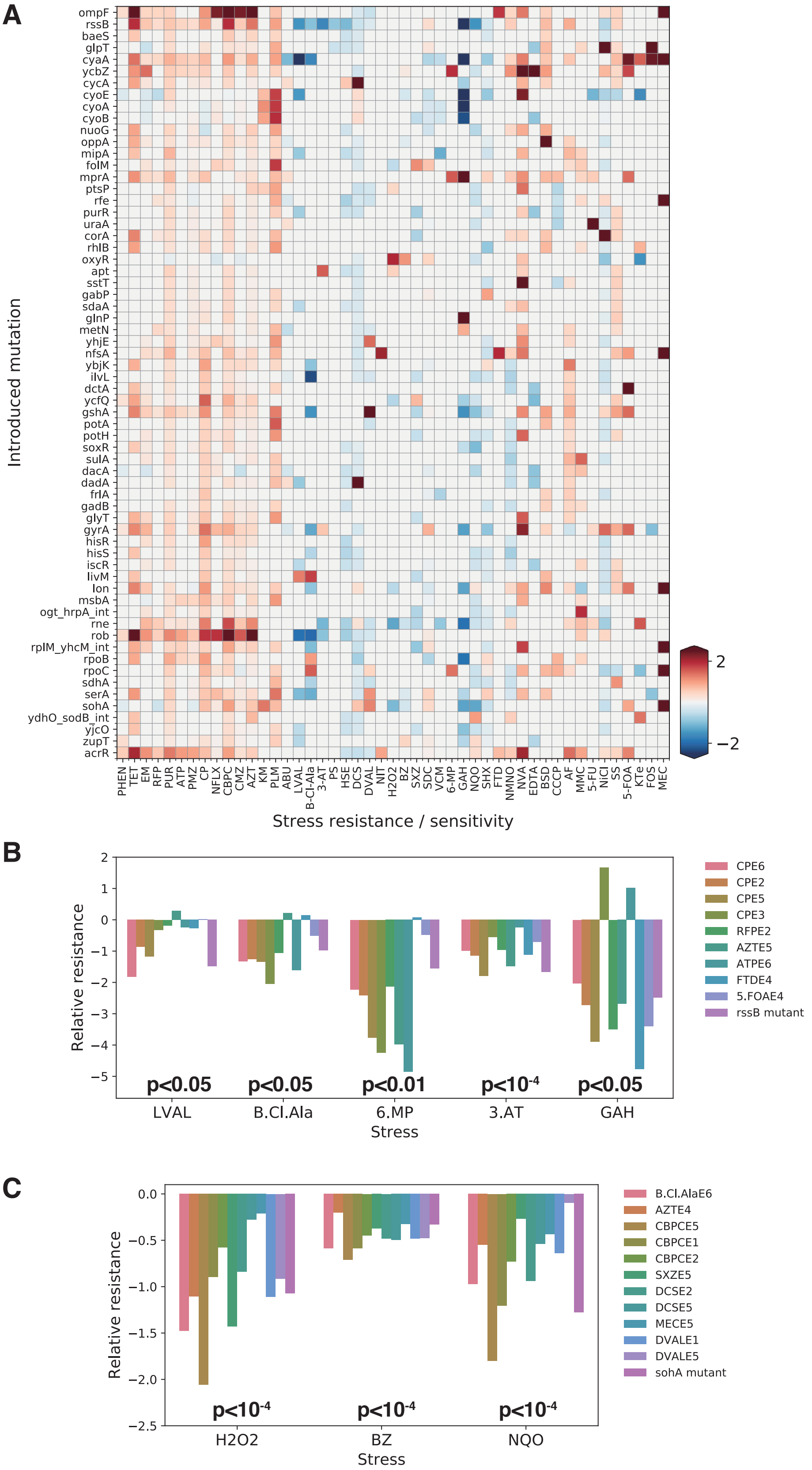


**Figure S4. Related to Figure 2 and Figure 3. Combinations of stresses which exhibited cross resistance, collateral sensitivity**

(A) Stresses which exhibited either resistance or sensitivity for the 64 reconstructed mutant strains (Mann-Whitney U-test, $p < 0.05$). Colors indicate the stress resistance relative to the parent strain. (B) and (C) Stress resistance relative to the parent strain for strains in class 9 and class 11, respectively. Resistance levels for the rssB and sohA mutant are also shown for comparison. The p-values were calculated by the Mann-Whitney U-test, which compared the evolved strains in the corresponding class and the parent strains.

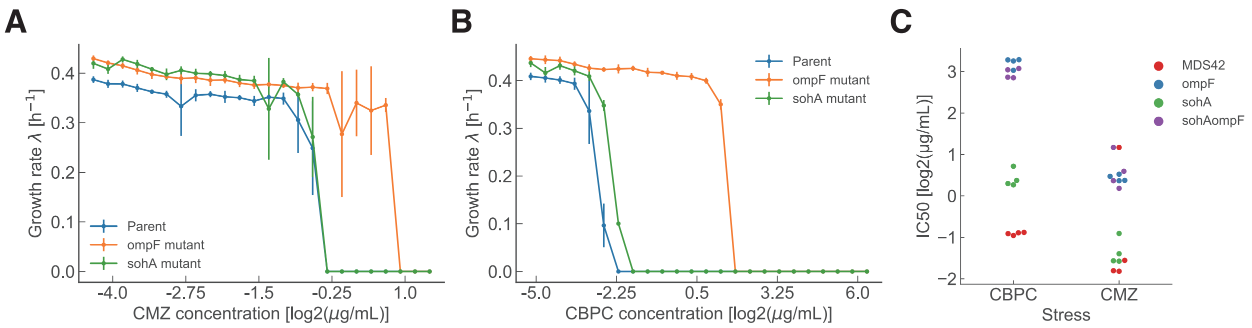


**Figure S5. Related to Figure 4. Neither fitness trade-offs nor epistasis explains decelerated evolution**

(A) and (B) Growth rates for the parent strain, ompF mutated strain, and the sohA mutant. Growth rates were measured in 24 concentration levels of cefmetazole (CMZ) and carbenicillin (CBPC), respectively. (C) IC50 levels measured for different stresses for the parent strain, ompF mutated strain, sohA mutated strain, and the sohA/ompF doubled mutated strain, respectively.
